# Tumor subclones, where are you?

**DOI:** 10.1101/2022.07.10.499466

**Authors:** Xianbin Su, Shihao Bai, Gangcai Xie, Yi Shi, Linan Zhao, Guoliang Yang, Futong Tian, Kun-Yan He, Lan Wang, Xiaolin Li, Qi Long, Ze-Guang Han

## Abstract

**Introduction:** Tumor clonal structure is closely related to future progression, which has been mainly investigated via mutation abundance clustering in bulk sample. With limited studies at single-cell resolution, a systematic comparison of the two approaches is still lacking.

**Methods:** Here, using bulk and single-cell mutational data from liver and colorectal cancers, we would like to check the possibility of obtaining accurate tumor clonal structures from bulk-level analysis. We checked whether co-mutations determined by single-cell analysis had corresponding bulk variant allele frequency (VAF) peaks. We examined VAF ranges for different groups of co-mutations, and also the possibility of discriminating them.

**Results:** While bulk analysis suggested absence of subclonal peaks and possibly neutral evolution in some cases, single-cell analysis identified co-existing subclones. The overlaps of bulk VAF ranges for co-mutations from different subclones made it difficult to separate them, even with other parameter introduced. The difference between mutation cluster and tumor subclone is accountable for the challenge in bulk clonal deconvolution, especially in case of branched evolution as shown in colorectal cancer.

**Conclusion:** Complex subclonal structures and dynamic evolution are hidden under the seemingly clonal neutral pattern at bulk level, suggesting single-cell analysis will be needed to avoid under-estimation of tumor heterogeneity.

**Research Highlights:**

- Bulk-level mutation abundance clusters are not equal to tumor subclones.
- Different groups of co-mutations could not be discriminated at bulk-level.
- Single-cell mutational analysis can identify rather than infer tumor subclones.
- Co-existing tumor subclones may have clonal neutral appearance at bulk-level.

**Lay summary:** Systematic comparison of tumor clonal structure differences between bulk and single-cell mutational analysis is lacking. Here we performed such as study and found that complex subclonal structures and dynamic evolution are hidden under clonal neutral appearance at bulk level in liver and colorectal cancers, suggesting single-cell analysis will be needed to avoid under-estimation of tumor heterogeneity.

## Introduction

Tumor is generally believed to be originated from mutations in a single cell, but when diagnosed the tumor mass usually contains large populations of progenies with different mutations and form subclones (Cairns, 1975; Nowell, 1976). The clonal structure and evolution within a tumor is closely related to its future progression such as treatment response and metastasis (Greaves and Maley, 2012; Marusyk et al., 2020; Yates and Campbell, 2012; Zahir et al., 2020). There are currently accumulative genomic data from bulk tumor tissues, providing insights on intra-tumor genetic heterogeneity (Dentro et al., 2021; Gerstung et al., 2020; Jamal-Hanjani et al., 2017; Turajlic et al., 2018). Many tools have also been developed to investigate tumor clonal structures based on the distribution of variant allele frequency (VAF) values from bulk samples, such as SciClone (Miller et al., 2014), PyClone (Roth et al., 2014) and MOBSTER (Caravagna et al., 2020a).

However, mutation cluster and tumor subclone are not equal items, and mutation co-occurrence is not available in bulk data but needs single-cell resolution confirmation (Gawad et al., 2014; Miles et al., 2020 ; Wang et al., 2014). Due to the high cost of single-cell DNA sequencing, most of single-cell studies focused on copy number alterations (Bian et al., 2018; Gao et al., 2017; Minussi et al., 2021 ; Navin et al., 2011), and there are only a few on tumor clonal structures from somatic mutations (Hou et al., 2012; Leung et al., 2017; McPherson et al., 2016). A systematic assessment of the difference of clonal structures from bulk and single-cell resolution analyses for the same tumor is essential to understand to what extent bulk data could depict genuine tumor subclones, but such a study is still lacking.

Here, we performed such a study by using both single-cell and bulk mutational data from liver and colorectal cancers, using both public datasets and newly generated data. We identified co-existing tumor subclones by single-cell mutational analysis, despite the absence of subclonal mutation clusters by bulk analysis. The results suggested that genuine tumor clonal structure may not be reliably revealed by bulk approach and will require single-cell dissection.

## Results

### Pseudo-bulk mutational analysis implied clonal neutral evolution in liver cancer

Inference of tumor subclones based on distribution of bulk-level VAF values is now widely used, and it is generally believed that the presence of mutation VAF clusters represents tumor subclones (Figure 1A). We have recently reconstructed single-variant resolution clonal evolution in liver cancer via patient-specific single-cell target sequencing (Su et al., 2021). As there were great inter-patient heterogeneities, in our previous work we used pseudo-bulk whole exome sequencing (WES) of single-cell genomic amplification mixture to screen for target mutations in each tumor. To better understand the pseudo-bulk mix WES, in this study we also generated true bulk WES data using the same specimens (HCC8-T, HCC8-PVTT, HCC9-T) for systematic comparisons (Figure 1B). The single-cell mutational profiles provided reliable clonal structure landscapes for cross-validation of bulk-level predictions.

**FIGURE 1.**
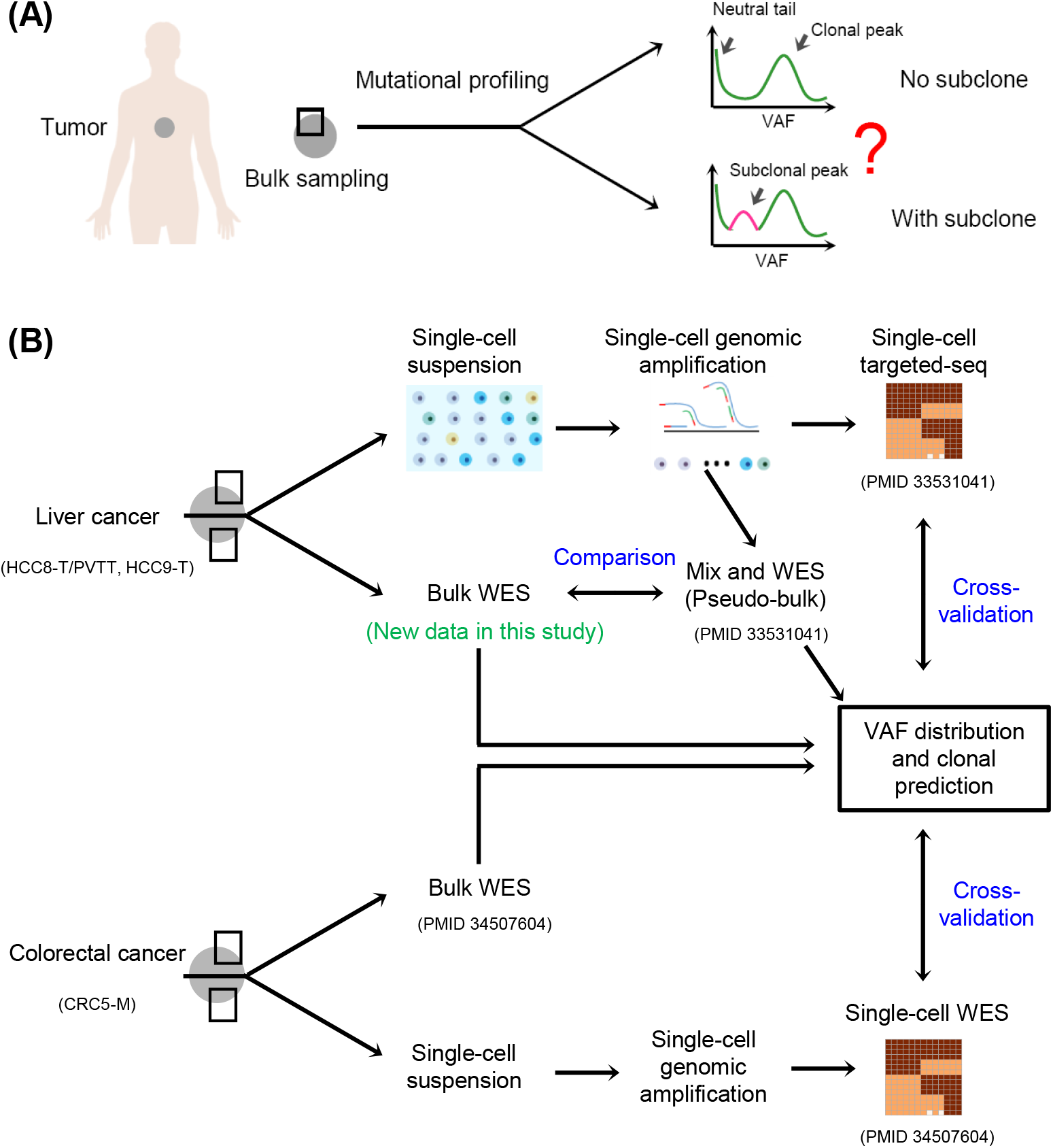
Overview of study design. (A) Schematic representation of tumor subclone inference via bulk-level mutational profiling. A typical bulk-level mutation variant allele frequency (VAF) distribution pattern includes a clonal peak and a neutral tail, and a subclonal peak between them is used as an indicator of the presence of a tumor subclone, which may be problematic. (B) Study design. Both liver and colorectal cancer bulk-level and single-cell mutational data were used for clonal structure analysis and cross-validation. For liver cancer, three samples were used (HCC8-T, HCC8-PVTT, HCC9-T), where HCC8-T and HCC8-PVTT are paired primary tumor and metastatic tumor thrombus from the same patient. Single-cell genomic amplification mixtures in liver cancer were used as pseudo-bulk samples for whole exome sequencing (WES), and single-cell targeted sequencing were used to get tumor clonal structures (Data from Su *et al*., *J Hematol Oncol* 2021, **14**(1):22, PMID 33531041). In this study we also generated new WES data using genuine bulk samples from the same liver cancer samples. For colorectal cancer sample CRC5-M, bulk WES data and single-cell WES data were used for mutation co-occurrence and VAF distribution analysis (Data from Tang *et al*., *Genome Med* 2021, **13**(1):148, PMID 34507604). Please note in both tumor types, different regions from the same tumor tissue were used separately for single-cell and bulk mutational profiling.

For the three liver cancer specimens, the distributions of VAF values from the mix approach exhibited similar pattern, with a clonal peak at VAF ∼0.5 and a cell division-related neutral tail containing mainly random mutations at VAF ∼0 (Figure 2A). It should be noted that the so-called neutral tail may contain low-frequency mutations that are related to future progression. Due to sequencing bias and allelic imbalance in bulk analysis, clonal mutations may span a wide VAF range, and the region between clonal peak and neutral tail is sometimes too narrow to discriminate subclonal VAF clusters. For liver cancer, there were no visible subclonal clusters between clonal peaks and neutral tails using two commonly used clonal analysis tools, SciClone and MOBSTER (Figure 2B,C), suggesting absence o f tumor subclones and possibly neutral evolution in all samples (Caravagna et al., 2020a; Williams et al., 2016; Williams et al., 2018).

**FIGURE 2.**
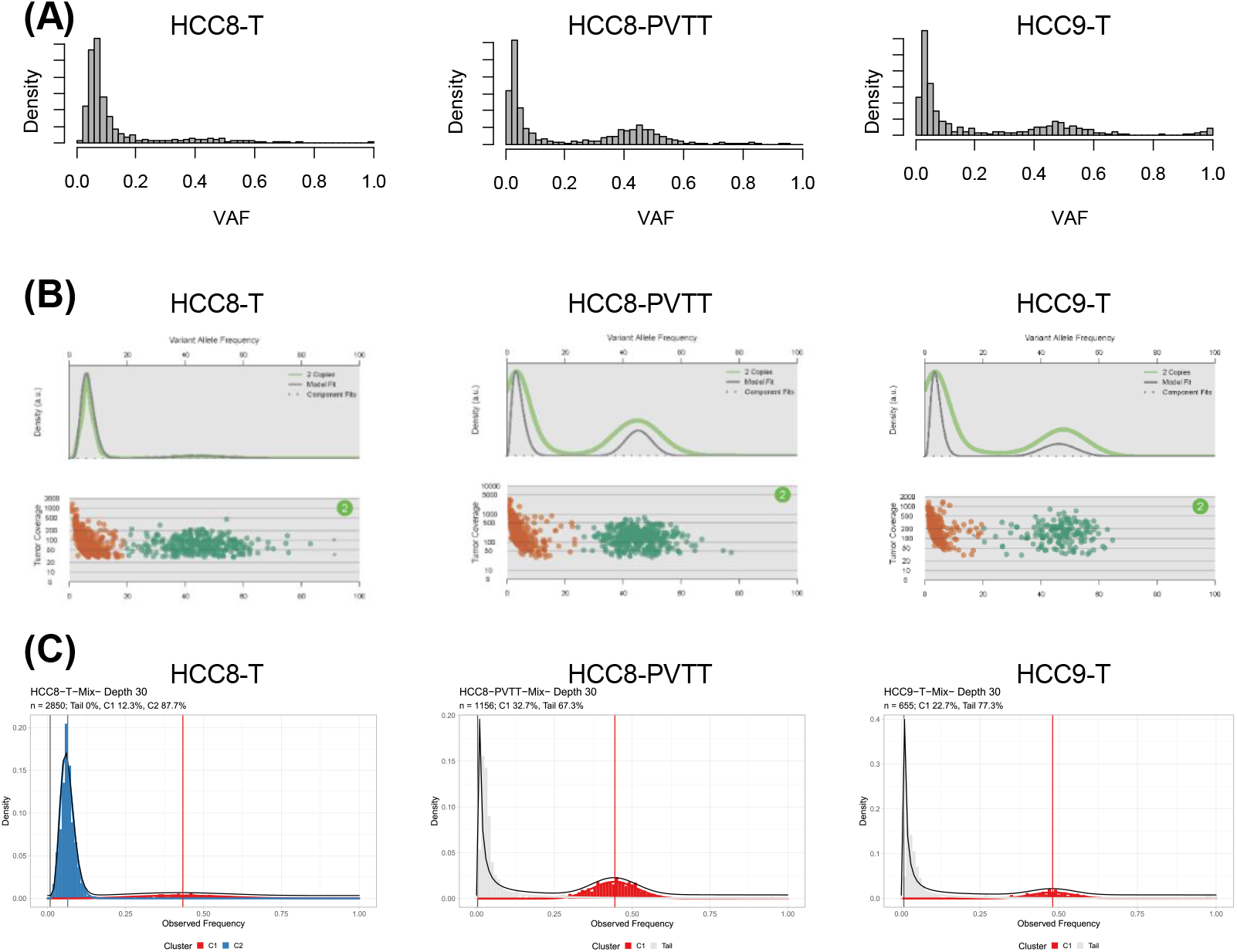
Pseudo-bulk mutational analysis implied absence of tumor subclones in liver cancer. (A) Distribution pattern of VAF values for mutations in three liver cancer samples derived from single-cell mix (pseudo-bulk) WES. (B-C) Subclonal deconvolution via SciClone (B) and MOBSTER (C) for the three liver cancer samples. Please note while Sciclone assigned the lower range VAF peak in each sample as a tumor subclone, MOBSTER recognized it as neutral tail in HCC8-PVTT and HCC9-T. The C2 cluster in MOBSTER result of HCC8-T should also be neutral tail.

### Single-cell analysis revealed co-existing tumor subclones in liver cancer

Single-cell target sequencing of somatic mutations, however, revealed a different scenario of clonal architectures in liver cancer. Co-existing subclones were identified by single-cell analysis in all samples, despite absence of subclonal clusters by bulk analysis. Three co-existing subclones with comparable sizes were identified in HCC8-T (Figure 3A), and the VAF ranges for clonal mutations and different groups of subclonal mutations had overlaps, suggesting that it may be difficult to assign a mutation to a specific subclone based on its bulk VAF value (Figure 3B). Similar results were found in HCC 8-PVTT and HCC9-T, with overlaps between different groups of co-mutations, implying that this is a general phenomenon in tumor clonal analysis.

**FIGURE 3.**
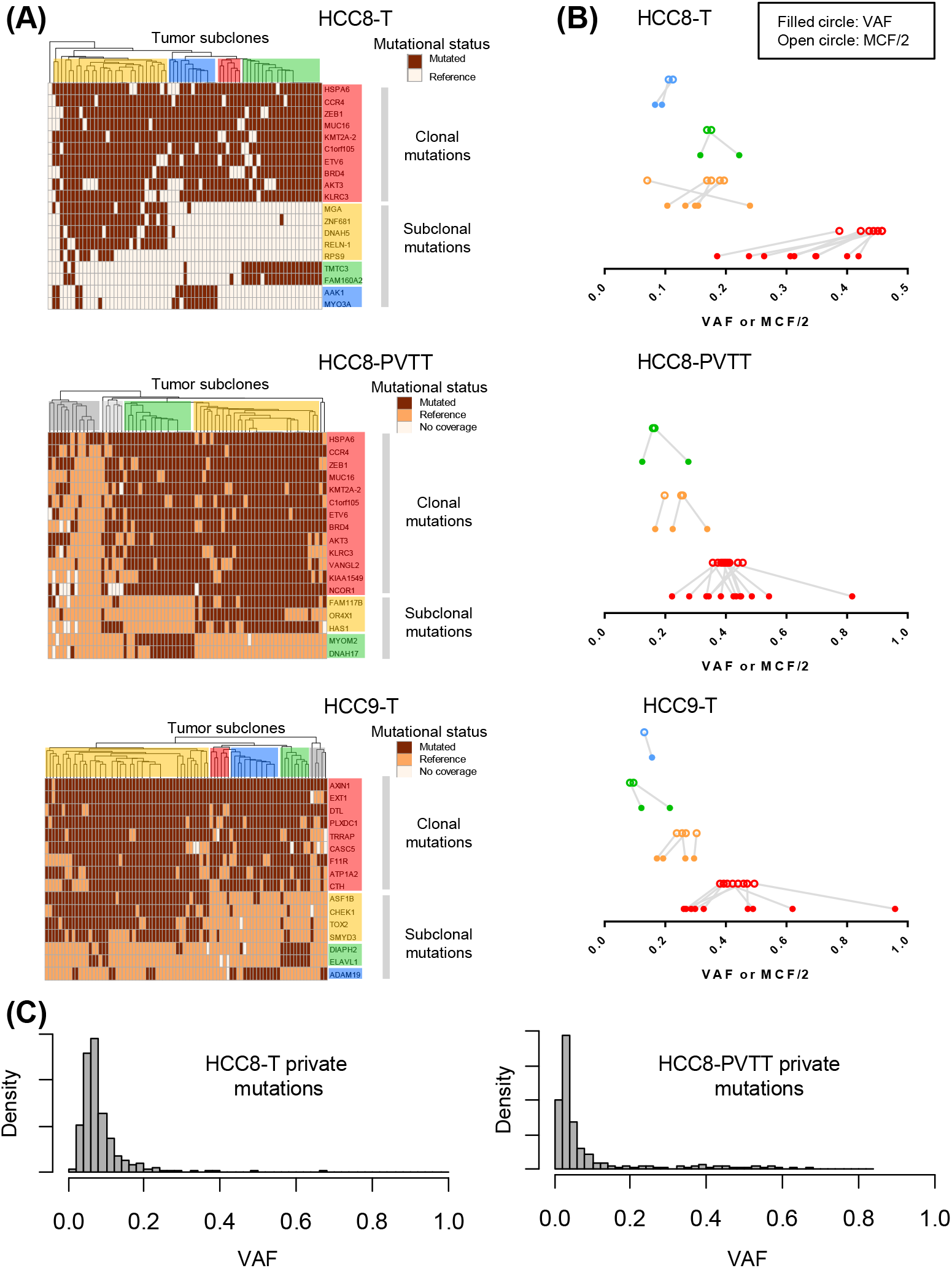
Single-cell analysis revealed co-existing tumor subclones in liver cancer not evident in mix approach. (A) Mutation co-occurrence in liver cancer samples revealed by single-cell analysis. Each row represented a somatic mutation and each column represented a cell. The color shading highlighted tumor subclones and corresponding mutations in each group. (B) Comparison of VAF and mutated cell fraction (MCF) values. To adjust for copy numbers, MCF/2 was used for comparison with VAF. Co-mutated clonal and subclonal mutations were grouped by single-cell analysis, with colors consistent with subclonal shading in (A). The lines indicated pairing VAF and MCF/2 for the same mutation. (C) VAF distribution of mutations privately found in HCC8-T or HCC8-PVTT.

Single-cell analysis also provided mutated cell fraction (MCF) value for each mutation, which is actually an indicator of subclone size. There were no overlaps between MCF ranges of clonal and subclonal mutations, although sometimes there were overlaps between MCF values from subclones with similar size (Figure 3B). A comparison of VAF and MCF showed that VAF generally had wider ranges in mix approach which may be more vulnerable to sequencing bias, making it difficult to infer clear clonal structures.

We then compared the mutations shared by or privately found in one specimens of HCC8-T and HCC8-PVTT, which were paired primary tumor and metastatic tumor thrombus from the same patient. Their shared mutations had higher VAF values, while their private mutations had relatively lower VAF values (Figure S1A). While shared mutations exhibited a clonal peak in both samples (Figure S1B), private mutations in each sample exhibited a neutral tail without detectable subclonal mutation cluster (Figure 3C). The results were consistent with previous finding of common origin and independent evolution for the two tumor specimens (Su et al., 2021). However, the absence of private subclonal clusters were contradicted by the presence of 3 and 2 subclones within each sample by single-cell analysis. A s the subclonal mutations in primary and metastatic tumors were not shared, they should not be introduced by early stage genetic drift but rather be acquired after occurrence of metastasis (Lynch et al., 2016). The results further supported that absence of subclonal VAF cluster in bulk analysis does not necessarily mean a lack of tumor subclone.

### No accurate tumor clonal structures in bulk-level analysis

As above tumor clonal analyses were conducted on pseudo-bulk single-cell mixtures, to rule out possible amplification bias or mixing imbalance, we then conducted genuine bulk WES on the same liver cancer samples. Most of mutations detected in the bulk approach were already found in the mix approach, and the latter also had more private mutations (Figure 4A). Different mutations in the two approaches could be attributed to tumor spatial heterogeneity, as they were actually profiling different regions of the same tumor sample (Sun et al., 2017). The shared mutations between two approaches also had higher VAF values while approach-private mutations had relatively lower VAF values (Figure S2A). Correlation analysis of shared mutations showed that VAF values from the bulk approach may be distorted by low tumor purity (Figure S2B).

**FIGURE 4.**
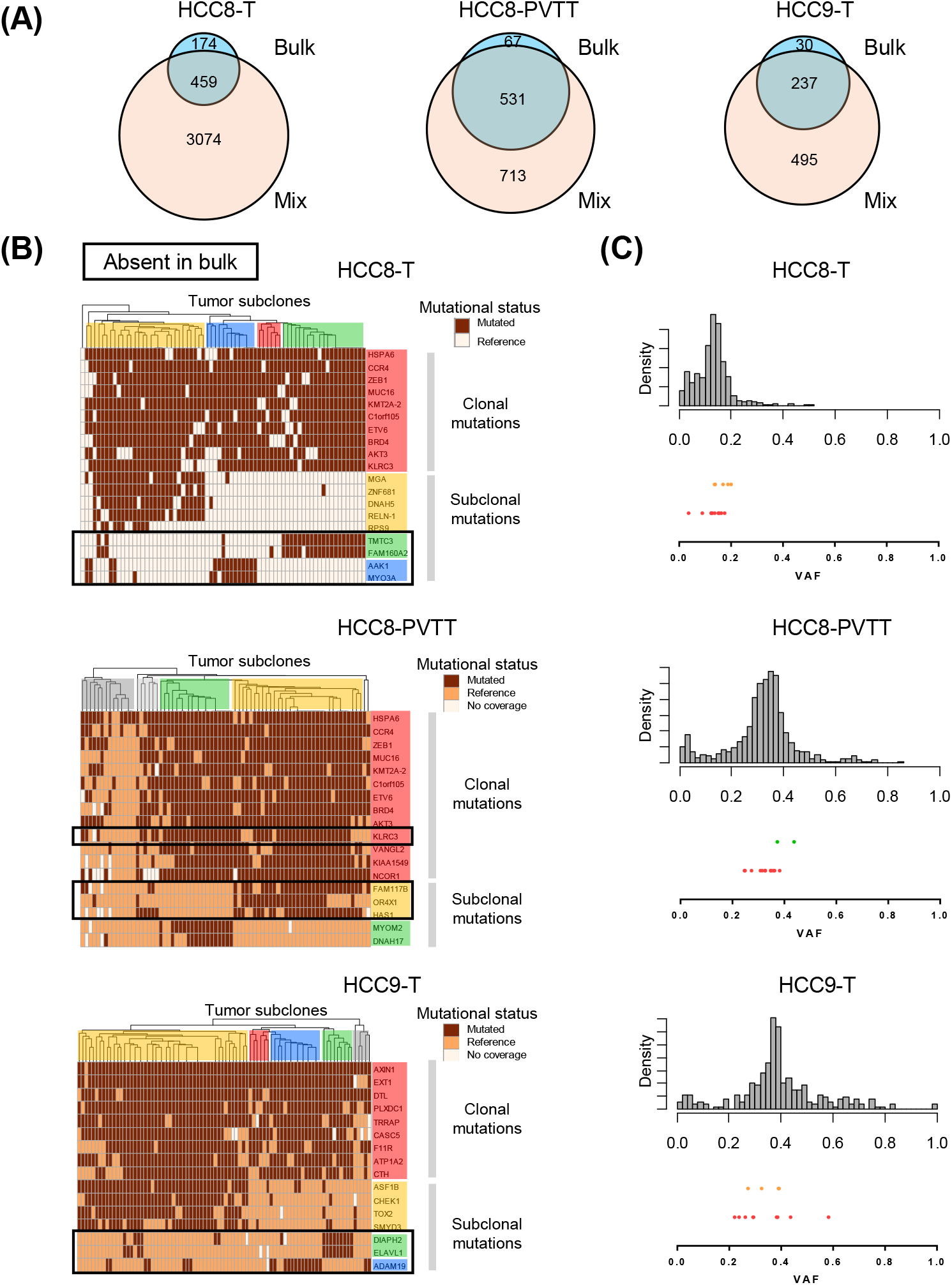
Accurate tumor clonal structures could not be revealed by bulk analysis. (A) Mutation overlaps between paired bulk and mix sequencing approaches in three liver cancer samples. (B) Mutations absent in bulk approach analysis shown in black boxes. (C) VAF distribution pattern of co-mutations in the bulk approach. The upper histogram showed VAF distribution of mutations from bulk-level WES, and lower part showed bulk VAF values for clonal and subclonal mutations grouped by single-cell analysis. Each dot represented a mutation, with colors consistent with subclonal shading.

For mutations included in single-cell target sequencing, there were subclonal mutation loss in all bulk samples, causing more simplified tumor clonal structures (Figure 4B). As for the recovered clonal and subclonal mutations, their VAF ranges also had overlaps, just as in the mix approach (Figure 4C). Considering tumor spatial heterogeneity, the results indicated that if bulk sample WES was used to guide downstream single-cell targeted mutational profiling, some subclones may be lost and the heterogeneities will be under-estimated. Besides single-cell mixture WES used in this study, WES using the same cell suspension for single-cell analysis (from the same tumor region) could be another reasonable choice which can be more relevant than neighboring tumor regions.

### Different groups of co-mutations could not be discriminated at bulk-level

We then checked whether different groups of co-mutations in liver cancer could be discriminated if other parameter was included besides VAF. As SciClone utilized depth *vs*. VAF for clonal analysis, we considered using ALT (number of altered reads). In the ALT *vs*. VAF plot, mutations in each sample formed two major clusters, with top right cluster representing mainly clonal mutations with higher VAF values and bottom left cluster representing neutral tail mutations with lower VAF values (Figure 5A). The region between the two clusters might contain subclonal mutations which may also overlap with the two clusters. As can be seen, co-mutations from single-cell analysis were intermingled together in the mix approach, making it difficult to separate them (Figure 5B). As there were no visible subclonal mutation clusters while single-cell analysis confirmed co-existing subclones, we concluded that accurate tumor clonal structures will require single-cell resolution dissection.

**FIGURE 5.**
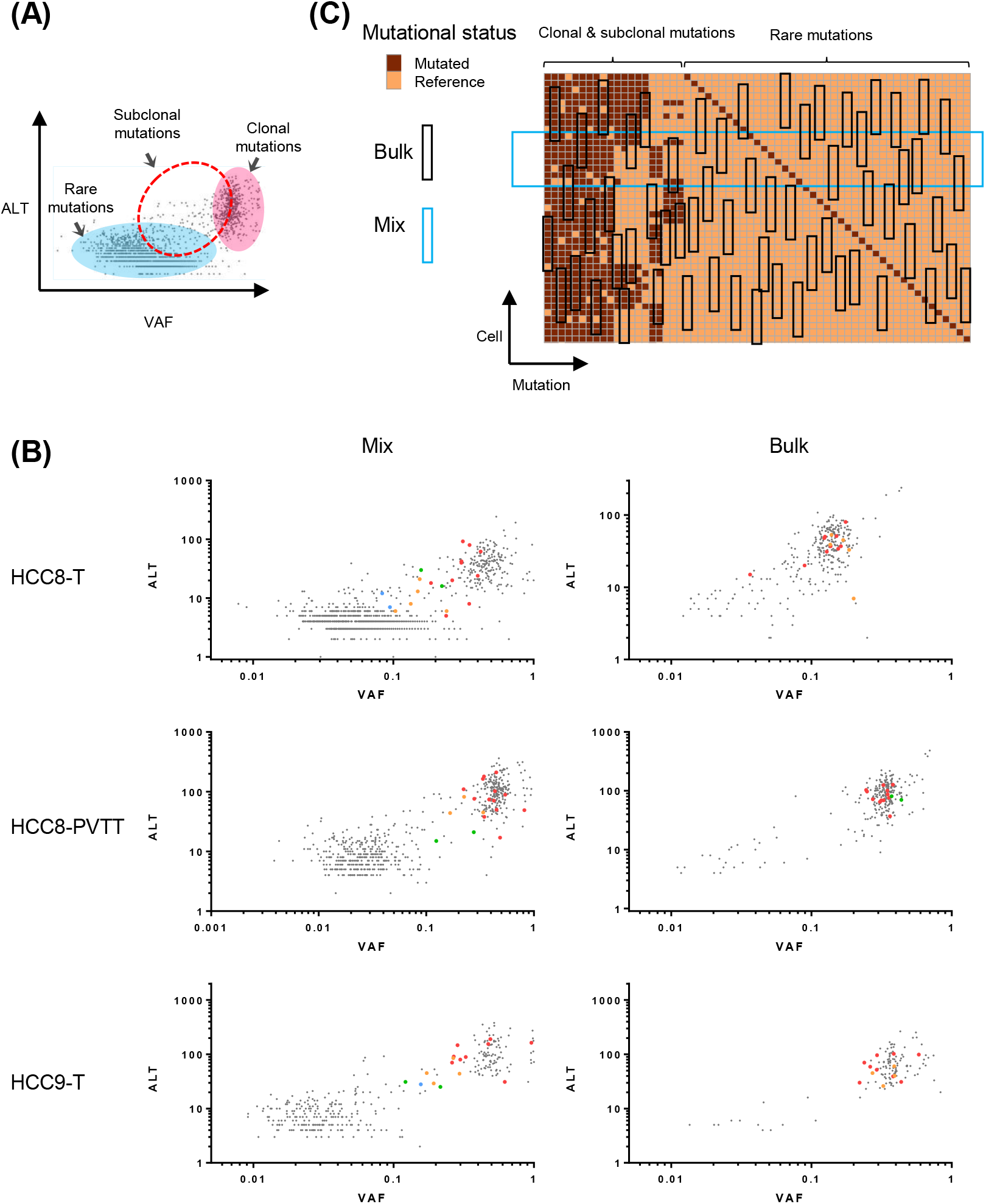
Different groups of co-mutations could not be discriminated in ALT *vs*. VAF plot. (A) Schematic representation of clonal, subclonal and rare mutations in ALT *vs*. VAF plot. (B) Distribution patterns of co-mutations in ALT *vs*. VAF plot in mix and bulk approaches. Color dots represented co-mutations from single-cell analysis, and grey dots represented other exonic mutations. (C) Sampling strategy difference between bulk and mix sequencing approaches. For single-cell mix approach, most clonal, subclonal and rare mutations are recovered as the sequencing coverage is at similar level with the number of single cells mixed (represented by the big blue box). For bulk approach, rare mutations will be easily lost due to random sampling at each mutation site (represented by the black box at each site).

In the ALT *vs*. VAF plot for the bulk approach, it was clear that there were less neutral tail mutations compared with th e mix approach (Figure 5B and S3), likely due to easier detection of rare mutations in a mixture from less than 100 single cells in comparison with random profiling more than millions of cells in the bulk approach (Figure 5C). Here the clonal and subclonal mutations were also intermingled, supporting that clear discrimination of co-mutations might be challenging in bulk approach, no matter from pseudo-bulk or genuine bulk samples.

### Dynamic evolution hidden under clonal neutral appearance at bulk-level

Single-cell WES has genomic coverage advantage in comparison with targeted sequencing, which will make clonal structure more reliable. As single-cell analyses in liver cancer were based on targeted sequencing, to rule out possible target selection bias or amplification distortion, we then analyzed a single-cell exonic mutational dataset from colorectal cancer (Tang et al., 2021). Sample CRC5-M exhibited branched evolution with step-by-step subclonal mutation acquisition and further split of each subclone (Figure 6A), demonstrating the complex relationship between subclonal mutation clusters and tumor subclones as the possession of a group of subclonal mutations may not always define a homogenous tumor subclone.

**FIGURE 6.**
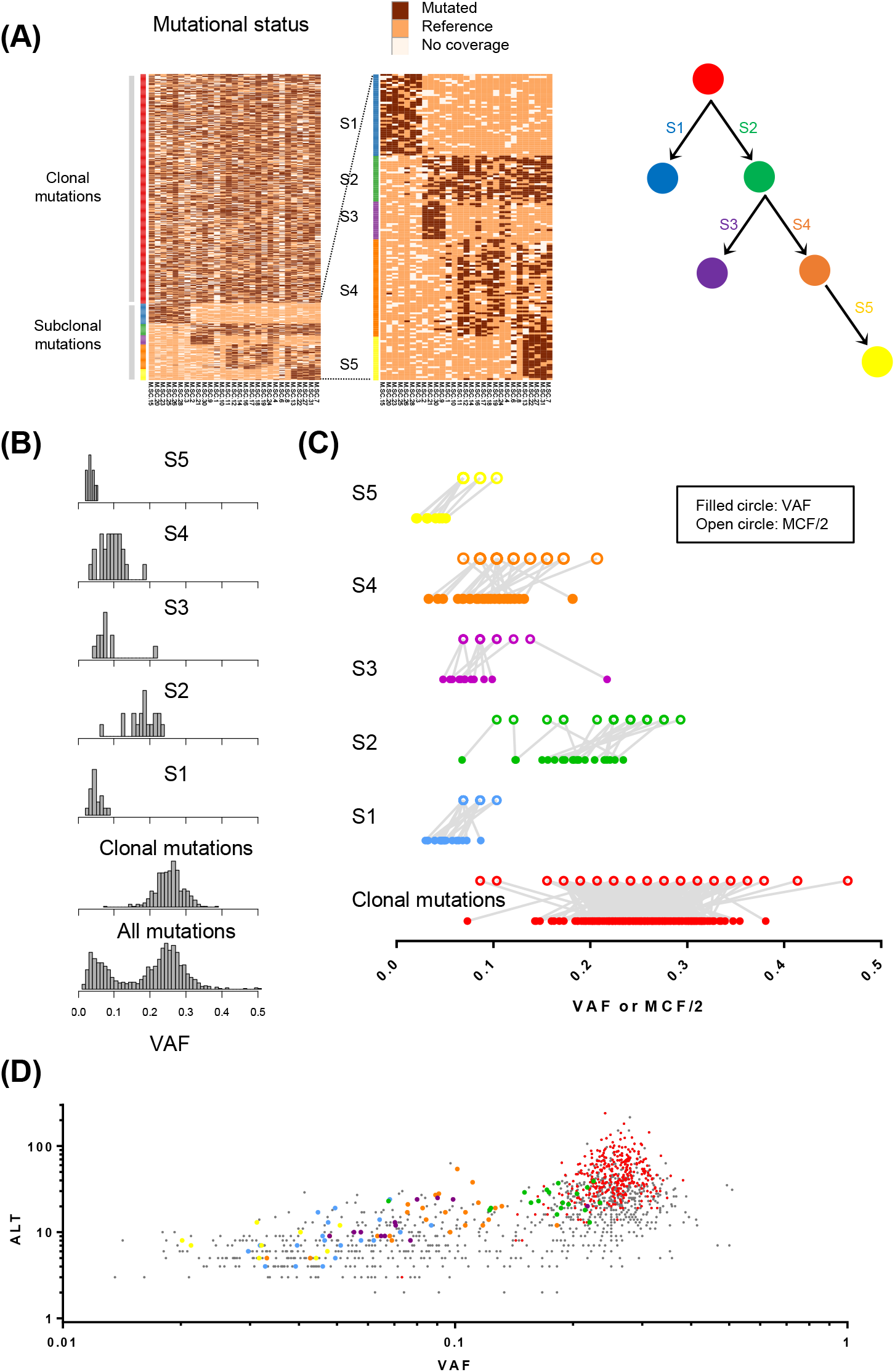
Dynamic evolution hidden under clonal neutral appearance at bulk-level in colorectal cancer. (A) Mutation co-occurrence in colorectal cancer sample CRC5-M revealed by single-cell WES. S1-S5 were subclonal mutation groups, and their acquisition order was shown on the right. (B) VAF distribution pattern of co-mutations. The histograms showed VAF distribution of different groups of subclonal mutations (S1-S5), clonal mutations, and all mutations from bulk-level WES. Please note the subclonal peaks were not reflected in the final histogram. (C) Comparison of VAF and MCF/2 values in co-mutated clonal and subclonal mutations, with colors consistent with subclonal shading in (A). The lines indicated pairing VAF and MCF/2 for the same mutation. (D) Distribution patterns of different groups of co-mutations in ALT *vs*. VAF plot.

Using the co-mutation groups defined by single-cell WES, we then checked their bulk VAF ranges in CRC5-M. Despite the complicated subclonal structure revealed by single-cell analysis, the VAF distribution showed a typical clonal peak and a neutral tail, and different groups of subclonal mutations were not reflected by corresponding subclonal peaks (Figure 6B). The VAF ranges of clonal mutations and Group S2 subclonal mutations had overlaps, while other groups of subclonal mutations (Group S1, S3, S4, S5) also had overlaps (Figure 6C). The ranges of VAF and MCF values for different co-mutation groups showed very good consistency in CRC5-M (Figure 6C), indicating reliable allelic representation in bulk exonic scale mutational profiling. In the ALT *vs*. VAF plot, the results also showed difficulty in discriminating different groups of co-mutations (Figure 6D). The analysis showed that tumor clonal structure was hidden under the seemingly clonal neutral pattern of bulk analysis, and accurate clonal structure and dynamic evolution will thus require investigation at single-cell resolution (Figure 7).

**FIGURE 7.**
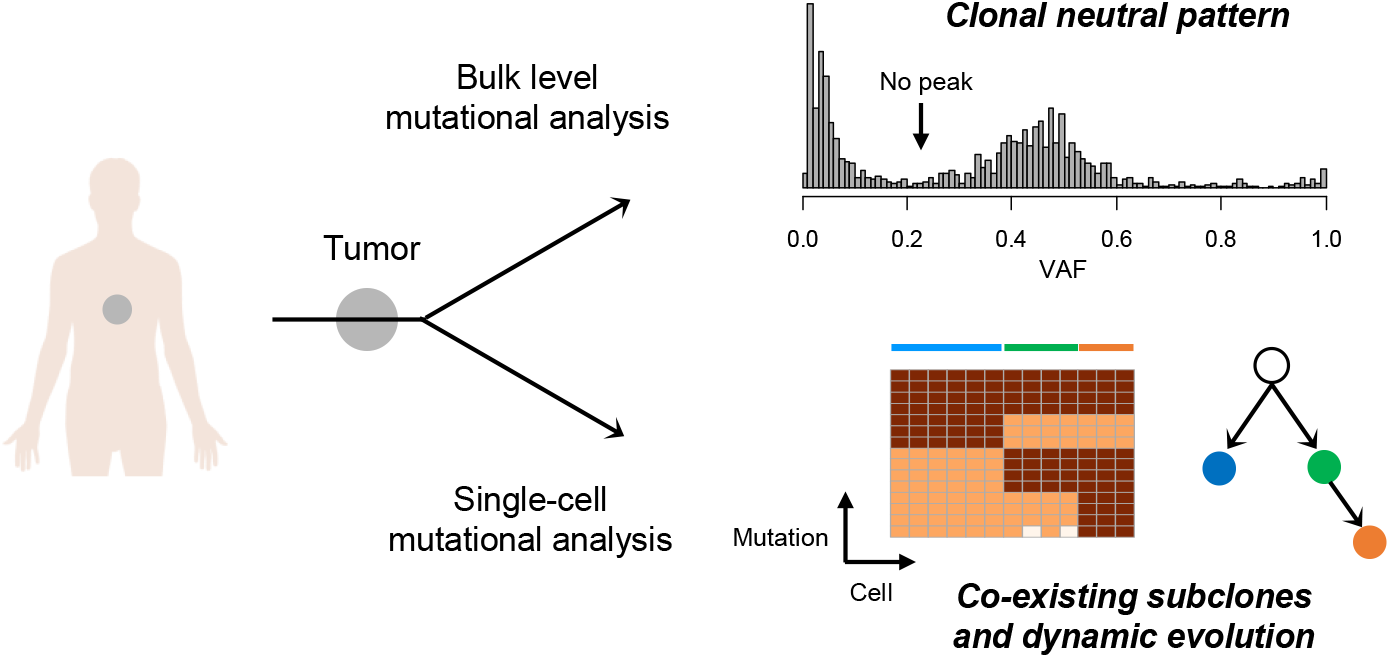
Schematic diagram showing the main finding. Complex tumor clonal structure and dynamic evolution could be revealed by single-cell analysis, which may be hidden under clonal neutral pattern in bulk analysis.

## Discussion

Bulk-level tumor clonal analysis has improved our understanding of intra-tumor heterogeneity, but it may not be able to reveal accurate tumor clonal architecture or reconstruct evolutionary history (Alves et al., 2017; Lim et al., 2020; Turajlic et al., 2019). Recently, a method that combined machine learning and population genetics was developed to enable more accurate subclonal reconstruction by ruling out interference from cell-division related neutral tail (Caravagna et al., 2020a). Ongoing subclonal selection was detected in 9 out of 298 high quality diploid tumor cases from PCAWG data, and prevalent neutral evolutionary pattern was proposed among tumors (Caravagna et al., 2020a). Here our analyses suggested that for some cases, the absence of subclonal mutation clusters does not necessarily support clonal neutral evolution, and utilization of such a criteria may underestimate the prevalence of tumor subclonal heterogeneity. Interpretation of clonal heterogeneity in bulk tumor samples should thus be careful, and systematic re-assessment of genetic heterogeneity in major tumor atlas datasets would be beneficial.

A major limit of bulk approach clonal analysis is the gap between mutation VAF cluster and tumor subclone, as they are two different terms that may not be exactly matched. For example, depending on the emerging stages of subclones during tumor progression, their subclone-specific mutations may not necessarily form apparent and detectable VAF clusters, especially for early subclones containing less mutations (Williams et al., 2018). Moreover, if there are subclones co-existing within a tumor at similar prevalence, their mutation VAF ranges will inevitably overlap and be difficult to separate. Tumor purity and genomic copy number status will further complicate the condition (Salcedo et al., 2020; Tarabichi et al., 2021), and clonal structure revelation thus calls for single-cell analysis (Davis et al., 2017; Evrony et al., 2021).

As it is difficult to obtain longitudinal specimens, dissection of clonal evolution is particularly challenging for solid tumors (Bailey et al., 2021). Single-cell profiling of tumor tissues based on somatic mutations will facilitate clonal history reconstruction at unprecedented accuracy, even for samples collected at a single time point (Dong et al., 2017; Evrony et al., 2021; Su et al., 2021). On account of still expensive whole genomic or exonic scale mutational analyses, however, the number of single cells profiled is still limited to hundreds for single-variant resolution studies (Duan et al., 2018; Tang et al., 2021; Wang et al., 2014). This will cause cell selection bias and lose rare subclones which might hold keys for treatment resistance or metastasis. Moreover, the spatial heterogeneity also makes it necessary to profile more than one region in a tumor by single-cell analysis, and this will need analysis of even more single cells. Current numbers of single cells analyzed were still a biased sampling of the tremendous genetic heterogeneities within tumors, and we expect future technological advances that enable mutational profiling of more single cells to shed light on tumor evolution and therapy design.

In summary, here we demonstrated that bulk-level analyses may be ill-suited for revealing tumor clonal structure due to difference between mutation cluster and tumor subclone. The absence of subclonal mutation cluster does not necessarily support clonal neutral evolution, and tumor clonal structure and evolution history can be better unveiled by single-cell analysis.

## Methods

### Clinical specimens and sequencing strategies of liver cancer

Single-cell mix (pseudo-bulk) WES and single-cell target mutation data from 3 liver cancer specimens were used in this study: HCC8-T, HCC8-PVTT and HCC9-T, in which HCC8-T and HCC8-PVTT were paired primary tumor and metastatic tumor thrombus from the same patient. Other samples with allelic dropout (ADO) issue or without subclones were not included in this study. Whole genome amplification product of single cells derived from paratumor and tumor tissues were separately mixed for WES, and ∼60 putative clonal and subclonal mutation sites were then selected from each patient for single-cell target sequencing (Su et al., 2021). The sequencing data for mix approach WES of HCC8-T, HCC8-PVTT and HCC9-T were obtained from project PRJNA606993 in NCBI SRA database, with BioSample accession number SAMN14118840, SAMN14118841 and SAMN14118843.

As a comparison between pseudo-bulk and genuine bulk approaches, the 3 liver cancer specimens also underwent bulk-level WES using Agilent SureSelect Human All Exon v7 K it (Agilent, 5191-4005) and illumina NovaSeq 2 × 150 bp sequencing mode. Sequencing reads were mapped to GRCh37/hg19 with BWA (Li and Durbin, 2009), mutations were called with GATK Mutect2 (McKenna et al., 2010), and SNPs were filtered using dbSNP141 (Sherry et al., 2001) and 1,000 Genomes Project (v3) database (Auton et al., 2015). The median sequencing depths for the tumor samples were more than 100×. The study was approved by the Ethnical Review Board of Shanghai Jiao Tong University, and the protocol conformed to the ethical guidelines of the 1975 Declaration of Helsinki.

### Subclonal deconvolution in liver cancer by mix approach sequencing

After calling mutations from each tumor sample, VAF values were calculated f or all mutations. Two tumor clonal analysis tools, SciClone (Miller et al., 2014) and MOBSTER (Caravagna et al., 2020a; Caravagna et al., 2020b), were then used to infer subclones in liver cancer samples. While SciClone separated the clonal peak (VAF ∼0.5) and neutral tail (VAF ∼0) but assigned both as subclones, MOBSTER could further recognize neutral tail in some cases.

### Tumor subclones and co-mutations in liver cancer single-cell data

Single-cell mutational data were used to investigate the clonal structures and mutation co-occurrence in each tumor case. After strict quality control, 71, 74 and 84 single cells from HCC8-T, HCC8-PVTT and HCC9-T were used for downstream analysis. Based on the mutational status of somatic mutations, single cells in each tumor were clustered into subclones. Clustering of mutations grouped them into clonal mutations present in all tumor cells, or subclonal co-mutations specifically found in each tumor subclone.

MCF for each mutation was calculated as the fraction of single cells harboring that mutation in a given tumor sample. Considering the effect of copy numbers, the ranges of VAF values in the mix approach and their corresponding MCF/2 values were compared to investigate the difference between mutation clusters and tumor subclones.

### Comparison of mutations in paired primary and metastatic liver tumors

For the paired HCC8-T and HCC8-PVTT, the plot of ALT *vs*. VAF (Shi et al., 2018) was used to check the VAF ranges of the shared and private mutations in each sample. The VAF distribution patterns of mutations shared by them or privately found in only one sample were compared to find possible subclonal peaks.

### Comparison of mutations in liver cancer by mix and bulk approaches

The numbers of shared and approach-private mutations were analyzed for the bulk and mix approaches. The plot of ALT *vs*. VAF was used to check the VAF ranges of the shared and private mutations in each approach. The correlation of VAF values between the two approaches for shared mutations were analyzed to check the extent of VAF deviation in different approaches, and recovery and loss of single-cell target mutations in the bulk approach were also analyzed to compare clonal structure difference. The VAF distribution patterns of mutations from the two approaches were shown in histogram plot. After clonal and subclonal mutations were grouped by single-cell analysis, VAF values of those grouped mutations in both mix and bulk approaches were compared to check range overlaps and relative locations to mutational peaks. The plot of ALT *vs*. VAF was also used to check the possibility of discriminating different groups of co-mutations in both bulk and mix approaches. The fractions of exonic mutation s with VAF <0.1 were calculated for comparison of neutral tail sizes in the mix and bulk sequencing approaches.

### Subclonal analysis of colorectal cancer

A recent work reported the clonal structure of both primary colorectal cancer and metastases based on single-cell WES, providing unbiased exonic scale single-cell mutational profiles (Tang et al., 2021). Here sample CRC5-M was chosen for subclonal analysis as it was t he sample with the most complicated subclonal structure and available bulk WES data. Mutation co-occurrences were revealed by single-cell analysis, and bulk VAF values of those co-mutations were compared to check range overlaps. A comparison of ranges of VAF and MCF/2 values for different groups of co-mutations were also performed. The plot of ALT *vs*. VAF was also used to check the possibility of discriminating different groups of co-mutations.

### Statistical analysis

Correlation analysis between two datasets were performed using GraphPad Prism 6, and R square values were provided for each analysis. Paired *t* test (Two-tailed) was performed to check the statistical significance between neutral tail sizes in the bulk and mix sequencing approaches in liver cancer.

## Data and materials availability

The sequencing data for bulk approach WES have been deposited in NCBI SRA database under project PRJNA606993, with BioSample accession number SAMN21591192, SAMN21591193 and SAMN21591194 for HCC8-T, HCC8-PVTT and HCC9-T. All other relevant data are available upon request.

## Acknowledgement

We would like to thank Professor Dan Xie and Dr. Kailing Tu from West China Hospital, Sichuan University for help with the single-cell WES data of colorectal cancer. This work was supported in part by National Natural Science Foundation of China (81802806, 31900484, 81902561), Guangdong Province Science and Technology Program (2020A1515010919), Natural Science Foundation of Shanghai (20511101900, 20ZR1427200), Shanghai Pujiang Program (20PJ1409800), SJTU Scientific and Technological Innovation Funds (2019TPA09) and SJTU Interdisciplinary Program (YG2021QN80).

## Author contributions

**X. Su:** Conception and design, methodology, formal analysis, investigation, data curation, writing - original draft, supervision, project administration. **S. Bai:** Formal analysis, writing - review & editing. **G. Xie:** Formal analysis, writing - review & editing. **Y. Shi:** Formal analysis, writing - review & editing. **L. Zhao:** Methodology, data curation. **G. Yang:** Data curation. **F. Tian:** Writing - review & editing. **K. He**: Investigation. **L. Wang**: Investigation. **Q. Long**: Conception and design, supervision, writing - review & editing. **Z. Han**: Conception and design, supervision, writing - original draft.

---

**FIGURE S1.**
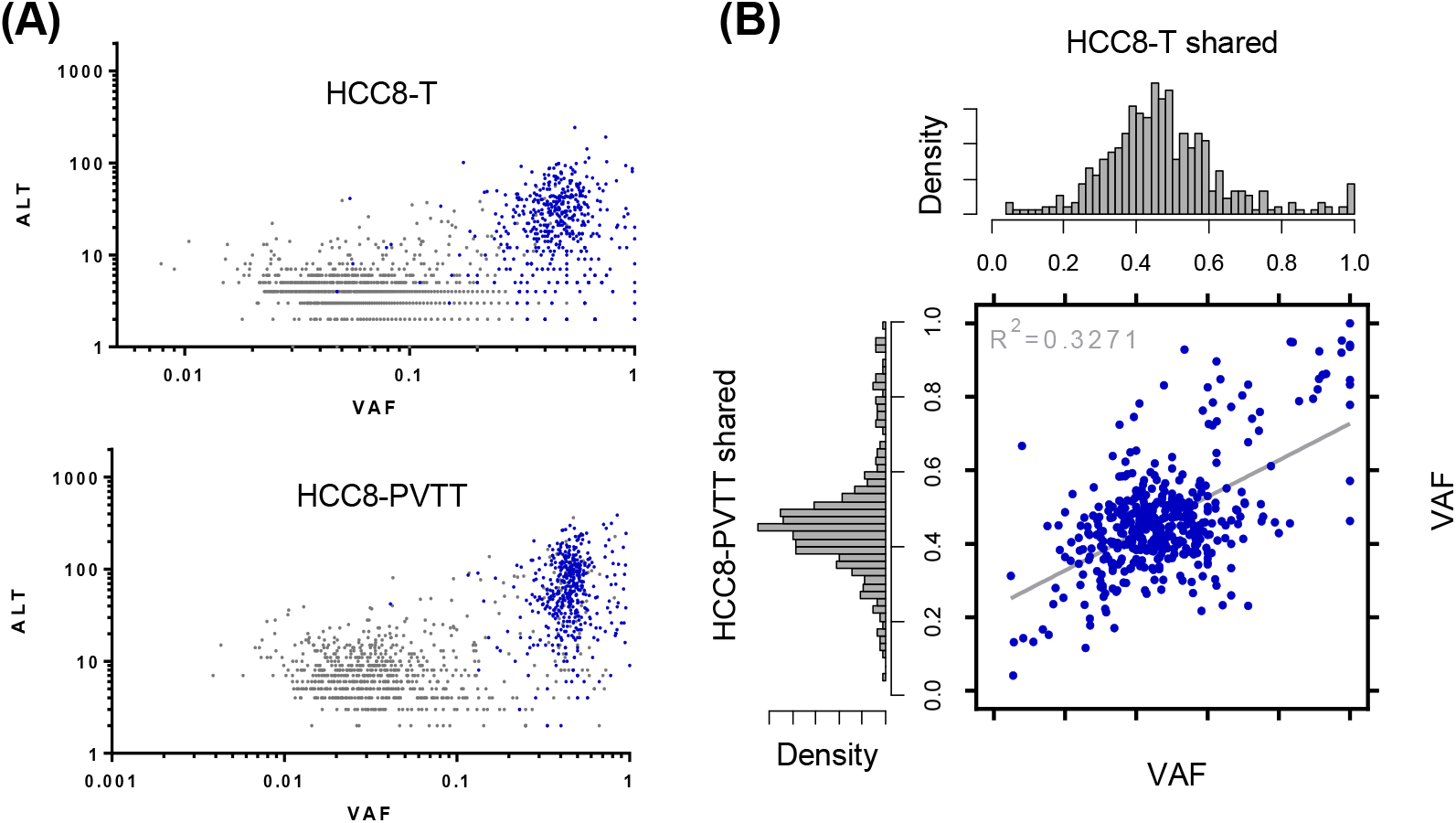
VAF distribution patterns of mutations from paired liver cancer samples. (A) Mutation overlaps between paired HCC8-T and HCC8-PVTT. ALT represented numbers of altered reads. Blue dots represented shared mutations, and grey dots represented sample-private mutations. (B) Correlation between VAF values in HCC8-T and HCC8-PVTT for shared mutations, with distribution histogram shown on the top and left for each sample.

**FIGURE S2.**
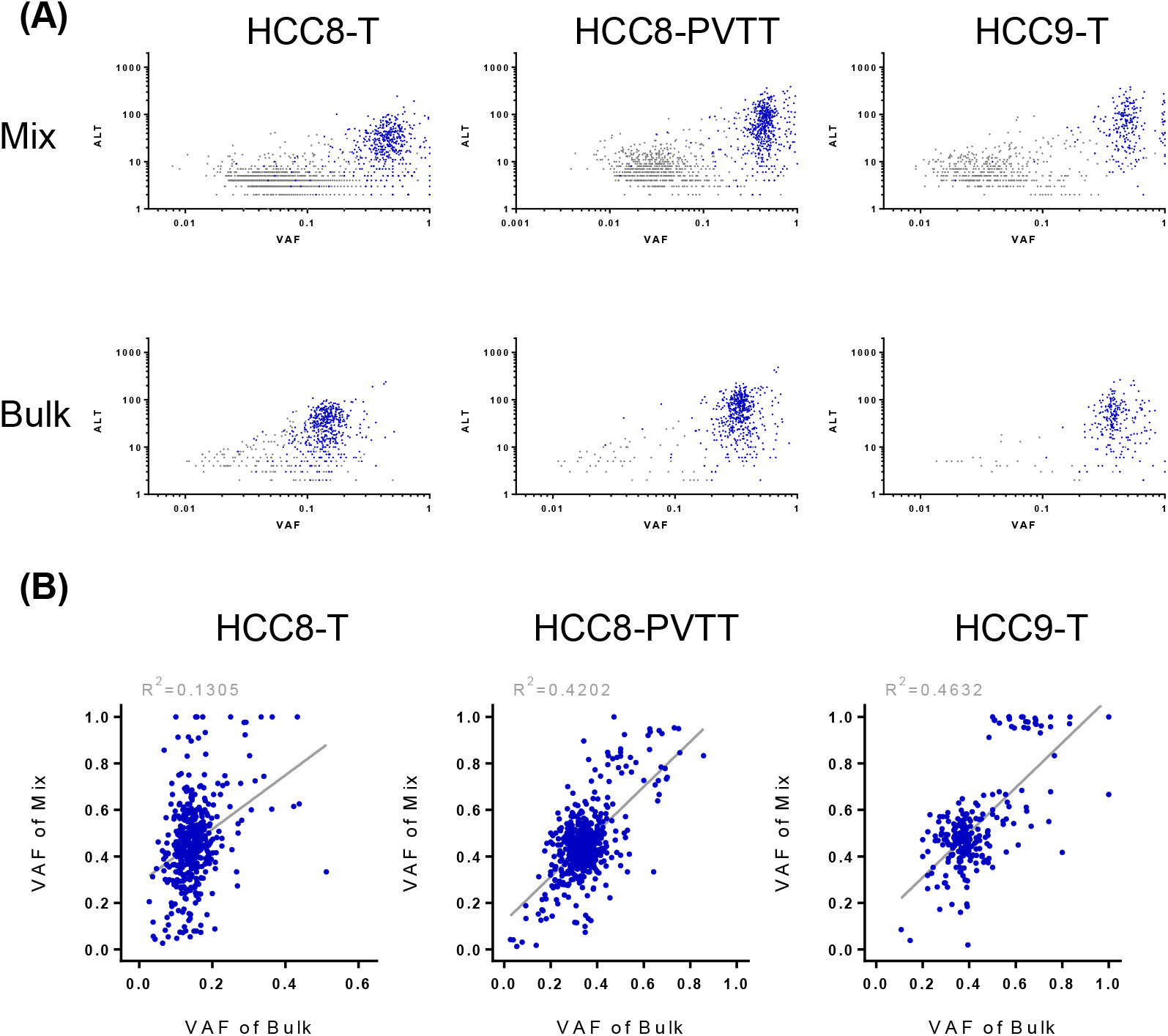
Mutation overlaps between bulk and mix sequencing approaches. (A) Mutation overlaps in ALT *vs*. VAF plot. Blue dots represented shared mutations, and grey dots represented private mutations in each approach. (B) Correlation between VAF values of shared mutations in bulk and mix approaches. The low R^2^ value in HCC8-T was caused by low tumor purity in bulk sample.

**FIGURE S3.**
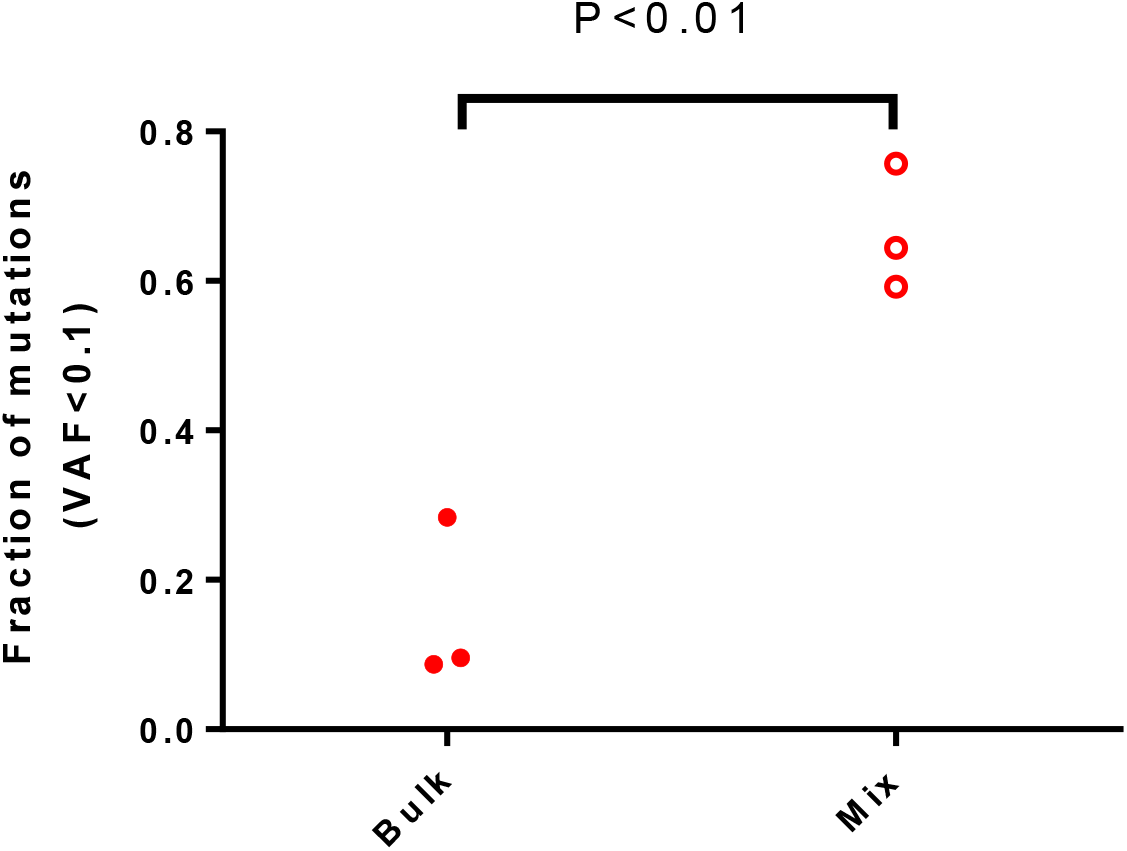
Comparison of neutral tail size in the bulk and mix sequencing approaches. The fraction of exonic mutations with VAF<0.1 was used as indicator of the neutral tail size for HCC8-T, HCC8-PVTT and HCC9-T. Paired *t* test (Two-tailed) was performed to check the statistical significance between the bulk and mix approaches.

